# Investigation of Gluthatione S-Transferase Variants in a Healht Population in Goiânia-Go

**DOI:** 10.1101/258806

**Authors:** Lucas Carlos Gomes Pereira, Nádia Aparecida Bérgamo, Angela Adamski da Silva Reis, Carlos Eduardo Anunciação, Elisangela de Paula Silveira-Lacerda

**Affiliations:** Laboratóirio de Genética Molecular e Citogenética, Instituto de Ciências Biológicas, Universidade Federal de Goiás, CEP: 74690-900, Goiânia/GO, Brazil; Laboratório de Genética Molecular, Instituto de Ciências Biológicas, Universidade Federal de Goiás, CEP: 74690-900, Goiânia/GO, Brazil

**Keywords:** Glutathione S-transferase, Genetic polymorphism, health population

## Abstract

Genetic polymorphisms in glutathione S-transferases (GSTs) genes might influence the detoxification activities of the enzymes predisposing individuals to a lot of disiases. Owing to the presence of these genetic variants, inter-individual and ethnic differences in GSTs detoxification capacity have been observed in various populations. Therefore, the present study was performed to determine the prevalence *GSTM1*0/*0, GSTT1*0/*0* and *GSTP1* Ile105Val polymorphisms in 100 healthy individuals from Goiânia - GO. *GSTM1* and *GSTT1* polymorphisms were analyzed by a Multiplex-PCR approach, whereas *GSTP1* polymorphisms were examined by PCR-RFLP. The frequencies of *GSTM1* and *GSTT1 *0/*0* genotypes are 49% and 31%, respectively. The frequencies of *GSTP1* Ile/Ile, Ile/Val, and Val/Val genotypes were 40%, 53%, and 7%, respectively. The wild-type (Ile) and variant (Val) allele frequencies were 66.5% and 33.5%, respectively. The combined genotypes distribution of *GSTM1, GSTT1* and *GSTP1* polymorphisms showed 12 possible genotypes present in our population; seven of them have a frequency greater than 5%. The effect of combined genotypes of these *GSTs* polymorphisms is still unknown. These findings in healthy population, give us such more information for the future epidemiological and clinical studies. Using to examine the effect of these combinations in drugs metabolism and cancer predisposition, further largest group would be needed, since their frequencies are quite low. To our of *GSTs* polymorfisms, this is the first study indicating the frequencies of genetic polimorphisms of GST superfamily in a health population in a Goiania population.

## 1. Introduction

Individual inherited genetic differences related to polymorphism in detoxification enzymes could be an important factor not only in metabolism but also in predisposition to pathologic condition [1]. Functional genetic polymorphisms have been described for Glutathione-S-transferase *(GSTs)* genes, a superfamily of phase II metabolizing enzymes. GSTs catalyze the conjugation of reduced glutathione (GSH) to a wide variety of electrophilic compounds in order to make them more soluble enabling their elimination [2]. As a result of this detoxification activity, GSTs protect the cell from DNA damage, genomic instability and cancer development. In addition, as non-enzymatic proteins, GSTs can modulate signaling pathways that control cell proliferation, cell differentiation and apoptosis, among other processes [3,4]. Deletion polymorphisms of *GSTM1* and *GSTT1* genes and the single nucleotide polymorphism in *GSTP1* c.319A > G (rs1695; p.105Ile > Val) lead to the absence or reducion of the detoxification capacity of the enzyme. Differences in GSTs activity may modify the risk of cancer development and also may impact on the heterogeneous responses to toxic substances or specific therapies [2]. Moreover, *GST* polymorphisms are known to contribute to inter-individual and ethnic variability in the susceptibility to environmental risk factors, cancer predisposition and drug responsiveness. Several epidemiological studies evaluated the role of *GST* polymorphisms on CML susceptibility, but conflicting results have been achieved [5,6]. There are no data available on these polymorphisms in the Goiania health population. The aim of the present study was to determine the prevalence of *GST* polymorphisms in the health population in Goiânia, a city located in the center of the west of Brazil, and to compare the results with different populations described in the literature.

## 2. Methods

### 2.1. Subjects

This study was carried out in the Genetic Molecular and Citogenetic Laboratory of the Institute of Biological Science I at Federal University of Goias, Goiânia, Brazil. The study was performed according approved by the ethics committee of Federal University of Goias (approval CEP: 895.552). In the current survey, 100 patients (35M/65F) of both genders and aged 18-90 (M: 30.2) years old considered healthy and practicing physical activity 2 to 3 times a week, were enrolled after receiving their informed consent.

### 2.2. Sample collection and DNA analyzes

Peripheral blood (5mL) was collected in EDTA vacutainer tubes from all participating individuals after obtaining their written consent. Until the DNA extraction blood DNA collections was store a -20^o^C. Genomic DNA extraction was performed from whole blood using a Purelink Genomic DNA Kit *(Invitrogen, Life Technologies Inc.; USA)*and the concentration was measured using a NanoDrop™ 2000/2000c Spectrophotometers *(Thermo Scientific, USA)*.

### 2.3. Genotyping

*GSTM1* and *GSTT1* polymorphism analyses were performed by multiplex polymerase chain reaction (PCR Applied Biosystems Veriti 9902, 96 Well Thermal Cycler), previously suggested by Abdel-Rahma, et al., [7], and modified by Reis, et al., [8], with the ubiquitous RH92600 gene as an internal standard. This was carried out amplification using the following primers: *GSTT1* forward primer 5` - TTC CTT ACT GGT CCT CAC ATC TC - 3`; *GSTT1* reverse primer 5` - TCA CCG GAT CAT GGC CAG CA - 3`; *GSTM1* forward primer 5`- GAA CTC CCT GAA AAG CTA AAG C - 3`; and *GSTM1* reverse primer 5` - GTT GGG CTC AAA TAT ACG GTG G - 3`. Each 25μL PCR reaction contained 2.5μL of 10X reaction buffer (Tris-HCl 10mM pH 8.3 and KCl), 2mM MgCl_2_, 200μM each of deoxynucleoside triphosphates, 10ρmol of each primer, 1 unit of Platinum Taq DNA polymerase (Invitrogen, Carlsbad, CA, USA), and 100ng genomic DNA. Thermal cycling conditions for the PCRs were as follows: 15min at 95°C, followed by 30 cycles of 95°C for 2min, 60°C for 1min, and 72°C for 1min, with a final extension at 72°C for 10min. PCR products were visualized on a 2% agarose gel electrophoresis at 100V for 50 min. Them the spected was confirmed two amplification bands, 480bp bands for *GSTT1* and 215bp for *GSTM1* were obtained for the *GSTT1+/GSTM1+* genotype. GSTT1+/GSTM1- genotype showed one band of 480bp, and the GSTT1-/GSTM1+ genotype showed a band of 215bp. For the *GSTT1-/GSTM1-* genotype (designated as null genotype), no bands were obtained.

*GSTP1* polymorphism analyses were performed by RFLP-PCR and the presence of 176pb DNA fragment in the samples were made using polyacrylamide gel that was stained with silver solution 4g L^-1^. After confirmed the amplification of 176bp band in all samples, the PCR product was restricted with the use of Alw26I *(Synapse)* enzyme according to the manufacturer’s suggested protocol for subsequent genotyping. Thermocycling was 12 hours at 37 °C followed by 20 minutes at 65 °C. The Alw26I restriction enzyme recognizes the DNA the GTCTC codon site at which occurs the exchange of the nucleotide adenine for guanine (occurring replacing the isoleucine amino acid at valine amino acid), and acts by cutting the DNA into two fragments of 91 and 85 bp, when there is the polymorphism.

### 2.4. Statistical analysis

The allelic and genotypic frequencies of the population were calculated using software Excel and expressed as a percentage. Fisher’s exact test was performed using software *GraphPad prism* version 7.00 for Windows, GraphPad Software, California USA.

## 3. Results

### 3.1. GSTM1 and GSTT1 deletion polymorphisms

In the current study, the frequencies of three polymorphisms in *GSTs* genes on individuals in a health population in Goiânia have been studied. The genotype distributions and allele frequencies observed in this study are shown in **Table 1**. We couldn’t calculate the expected Hardy–Weinberg frequencies for *GSTM1* and *GSTT1* genotypes, as multiplex-PCR approach cannot differentiate between the wild-type (*1/*1) and heterozygous (*0/*1) genotypes. The *GSTM1* and *GSTT1* deletion polymorphisms were analyzed by Multiplex-PCR approach, using Rh92600 as an internal control. As shown in Table 1, the frequencies of *GSTM1* and *GSTT1 *0/*0* genotypes are 49% and 31%, respectively.

**Table 1.** Distribution of genotype and allele frequencies of GSTM1 and GSTT1, polymorphisms in a health population in Goiânia.

|  | Age |  |  | Gender |  | Total |
| --- | --- | --- | --- | --- | --- | --- |
|  | 18 - 30 | 31 – 50 | ≥ 51 | Male | Female |  |
| Number of Individuals | 62 | 33 | 5 | 35 | 65 | 100 |
| *GSTM1* null frequency (%) | 33 (0.33) | 14 (0.14) | 2 (0.02) | 15 (0.15) | 34 (0.34) | 49 (0.49) |
| *GSTT1* null frequency (%) | 20 (0.20) | 9 (0.09) | 2 (0.02) | 9 (0.09) | 22 (0.22) | 31 (0.31) |

### 3.2. GSTP1 Ile105Val polymorphism

The *GSTP1 Ile105Val* polymorphism was performed by PCR-RFLP. As shown in Table 2, the frequencies of *GSTP1 Ile/Ile, Ile/Val*, and *Val/Val* genotypes were 40%, 53%, and 7%, respectively. The wild-type *(Ile)* and variant *(Val)* allele frequencies were 66.5% and 33.5%, respectively (**table 2**).

**Table 2.** Genotype and allele frequencies of GSTP1 Ile105Val polymorphism of Tunisian population and various ethnic groups.

| Genotypes | N | Allele frequencies |  |
| --- | --- | --- | --- |
|  |  | ILE | VAL |
| *ILE/ILE* | 40 |  |  |
| *ILE/VAL* | 53 | 0,665 | 0,335 |
| *VAL/VAL* | 7 |  |  |

### 3.3. Combined genotype analysis

The distribution of combined genotype of *GSTM1, GSTT1* and *GSTP1* polymorphisms showed the possible genotypes are present in a health population from Goiânia. Among these, seven genotype combinations showed frequency greater than 5%. The most frequently observed combinations were null M1/ non-null T1/ Ile/Val (20%), non-null M1/ non-null T1/ Ile/Ile (15%), non-null M1/ non-null T1/ Ile/Val (15%) and null M1/ non-null T1/ Ile/Ile (14%) (**Table 3**).

**Table 3.** Combined genotype analysis of GSTM1, GSTT1 and GSTP1 polymorphisms.

| *GSTM* | *GSTT* | *GSTP* | Frequency |  |
| --- | --- | --- | --- | --- |
|  |  |  | n | % |
| Non Null | Non Null | *Ile/Ile* | 15 | (0.15) |
|  | Non Null | *Ile/Val* | 15 | (0.15) |
|  | Non Null | *Val/Val* | 3 | (0.03) |
|  | Null | *Ile/Ile* | 4 | (0.04) |
|  | Null | *Ile/Val* | 12 | (0.12) |
|  | Null | *Val/Val* | 2 | (0.02) |
| Null | Non Null | *Ile/Ile* | 14 | (0.14) |
|  | Non Null | *Ile/Val* | 20 | (0.20) |
|  | Non Null | *Val/Val* | 1 | (0.01) |
|  | Null | *Ile/Ile* | 7 | (0.07) |
|  | Null | *Ile/Val* | 6 | (0.06) |
|  | Null | *Val/Val* | 1 | (0.01) |
| Total |  |  | 100 | 100% |

n: number of subjects.
Combined genotypes were expressed by percentage (%).
Null: homozygous deletion (\*0/\*0).
Non-null: heterozygous deletion (\*0/\*1) or wild homozygous (\*1/\*1).

## 4. Discussion

The result for *GSTM1* and *GSTT1* deletion polymorphisms concerning the *GSTM1*0/*0* and *GSTT1*0/*0* genotype was 49% and 31%. This is an agreement with previous report of Kubiszeski *et al*. [9] in Cuiabá population (50% and 32%), and Rocha *et al*. [10] in Amazônia population. Other studies in the Brazilian population from Bahia, performed by Rocha *et al*. [10], and Pinhel, *et al*. [11] in the population from São Paulo confirm our findings in the Goiânia population. *GSTM1* and *GSTT1* enzymes metabolize several precarcinogens, drugs, constituents of tobacco smoke and solvents to reactive metabolites which ultimately lead to DNA or protein damage [12]. Hence, the data on the prevalence of this polymorphism will help in predicting susceptibility to various cancers.

The results for *GSTP1 Ile105Val* polymorphism are similar to those reported in São Paulo population (n 53), in which the wild-type (Ile) and variant (Val) allele frequencies were 67% and 33% (P=1,000), respectively [13] and in Rio de Janeiro population (n 531) with 69% and 31% (P=0,7628) respectively [14], both studies with health people.

Regarding to the other ethnic groups, Goiânia population showed a similar prevalence of *GSTP1* Ile105Val with those reported in some Asian populations such as, Chinese, Indian, Tunisian and venezuelan [12, 15, 16, 17, 18, 19] as well as with White Americans and South Africans [20, 21]. Interestingly, Goiânia population showed a significantly high prevalence of *GSTP1* Ile105Val polymorphism than that reported in Japanese population, in which the wild-type (Ile) and variant (Val) allele frequencies were 84% and 16% (P=0,0081*), respectively [21] and the Korean population with 80,9% and 19,1%, respectively (P= 0,0355*) [22].

The importance of this class of enzymes and, in a general way, of all the enzymes of the GST family has been increasingly highlighted by its relation in several processes of metabolism and catalytic properties and of detoxification besides having discovered other biologically important functions as, for example, in protein-protein interactions; involvement with chaperones and mechanisms of kinases; and especially myeloproliferative properties [23].

*GSTP1* is a major enzyme metabolizing anti cancer drugs like oxiplatin, cyclophosphamide which are used in the treatment of breast cancer and colorectal cancer [24]. An over expression of this enzyme in individuals with Ile/Ile genotype causes resistance to drugs like cisplatin [25, 26]. Therefore, investigation of this polymorphism will provide a clue to the identification of responders to cancer therapy with certain chemotherapeutic drugs.

Regarding the effects of the combined genotypes, some published reports showed that single *GST* gene polymorphism does not significantly increase risk to cancer [27], suggesting that investigations on combined genotypes of *GSTM1, GSTT1* and *GSTP1*, or even in relation to other metabolizing enzymes are needed. Additionally, some other studies have reported a relationship between the combination of *GSTM1, GSTT1*, and *GSTP1* genotypes and the risk of various diseases, such as chronic lymphocytic leukemia, thyroid cancer and they suggested a possible synergistic effect between *GST* genotypes [28, 29].

Genetic polymorphisms of *GSTs* genes differ significantly among racial groups and residential populations in different parts of the world [30, 31]. Based on our findings, Goiânia population showed an, eventual, similarity with others Brazilian populations for genotype and allele frequencies of *GSTM1, GSTT1, GSTP1* polymorphisms.

The obstacle with research using the Brazilian population is that it resulted basically from the racial mixture between whites from the Iberian Peninsula and Africans of various ethnical groups, with a small contribution by native Amerindians. However, this may also represent the lack of a rigid distinction between races and the intense admixture that has been occurring in this country [32].

Studies, including the present one, indicate that while evaluating the role of a particular *GST* gene in any disease susceptibility, the whole pattern of different biotransformation enzymes should be considered. Wormhoudt *et al.*,[33] explains this is because multiple detoxification enzymes may be involved in the metabolism of a given compound and the resulting metabolites may produce different effects clinically.

The effect of combined genotypes of these *GSTs* polymorphisms is still unknown. These findings in healthy population, give us such more information for the future epidemiological and clinical studies. Using to examine the effect of these combinations in drugs metabolism and cancer predisposition, further largest group would be needed, since their frequencies are quite low.

## 5. Conclusion

This study provides the first results of genotype distribution and allele frequencies of *GSTs M1, T1* and *P1* polymorphisms in a health population in Goiânia – GO. An identifier of polymorphisms related to predisposition may contribute to the implementation of a public health policy focused on preventive medicine. Hence, it opens up new avenues for further investigations by epidemiologists in determining inter individual variation in genetic susceptibility to various diseases caused due to gene-environment interaction.

